# Generation of light-producing somatic-transgenic mice using adeno-associated virus vectors

**DOI:** 10.1101/328310

**Authors:** Rajvinder Karda, Ahad A. Rahim, Andrew M.S. Wong, Natalie Suff, Juan Antinao Diaz, Dany P. Perocheau, Nuria Palomar Martin, Michael Hughes, Juliette M.K.M. Delhove, John R. Counsell, Jonathan D. Cooper, Els Henckaerts, Tristan R. Mckay, Suzanne M.K. Buckley, Simon N. Waddington

## Abstract

We have previously designed a library of lentiviral vectors to generate somatic-transgenic rodents to monitor signalling pathways in diseased organs using whole-body bioluminescence imaging, in conscious, freely moving rodents. We have now expanded this technology to adeno-associated viral vectors. We first explored bio-distribution by assessing GFP expression after neonatal intravenous delivery of AAV8. We observed widespread gene expression in, central and peripheral nervous system, liver, kidney and skeletal muscle. Next, we selected a constitutive SFFV promoter and NFκB binding sequence for bioluminescence and biosensor evaluation.

An intravenous injection of AAV8 containing firefly luciferase and eGFP under transcriptional control of either element resulted in strong and persistent widespread luciferase expression. A single dose of LPS-induced a 10-fold increase in luciferase expression in AAV8-NFκB mice and immunohistochemistry revealed GFP expression in cells of astrocytic and neuronal morphology. Importantly, whole-body bioluminescence persisted up to 240 days.

To further restrict biosensor activity to the CNS, we performed intracerebroventricular injection of each vector. We observed greater restriction of bioluminescence to the head and spine with both vectors. Immunohistochemistry revealed strongest expression in cells of neuronal morphology. LPS administration stimulated a 4-fold increase over baseline bioluminescence.

We have validated a novel biosensor technology in an AAV system by using an NFκB response element and revealed its potential to monitor signalling pathway in a non-invasive manner using a model of LPS-induced inflammation. This technology employs the 3R’s of biomedical animal research, complements existing germline-transgenic models and may be applicable to other rodent disease models with the use of different response elements.

## Introduction

Germline light producing transgenic mice where luciferase expression is controlled by an endogenous promoter, a surrogate promoter or by a minimal promoter downstream of tandem, synthetic, transcription factor binding elements, are used to provide an *in vivo* readout of physiological and pathological processes^1,2^. One of the advantages of this technology is that every cell will contain a copy of the luciferase transgene and therefore provide a whole-body transgene expression profile under the control of a specific promoter of choice. However, producing germline transgenics requires frequent backcrossing and therefore becomes a time-consuming and costly process, using many rodents.

We have previously developed a novel technology which allows the generation of light producing somatic transgenic rodents, using lentiviral vector as a proof-of-concept system^3^ and have validated this technology both *in vitro*^4,5^ and *in vivo*^1,6^. We have also demonstrated that signalling pathways in diseased organs can be monitored specifically, continually and in a non-invasive manner^1,2^. Exploiting the immune tolerance of neonatal mice towards transgenic proteins^7^, we were able to achieve organ specific transduction by a single neonatal administration of the biosensor.

To achieve a better spread of delivery and target additional tissues, we explored adeno-associated viral (AAV) vectors to deliver the biosensor construct. Previous work has shown that AAV8 can achieve widespread transgene expression and cross the blood brain barrier (BBB) after an intravenous administration in adult rodents, targeting the brain, heart, liver and skeletal muscles^8^. Neonatal intraperitoneal administration of an AAV8 vector resulted in expression in pancreas^9^, kidney^10^ and spinal cord^11^.

In this study, we have shown extensive, systemic gene expression after a single neonatal intravenous administration of AAV8-CMV-eGFP to neonatal mice. We have generated AAV8 biosensors carrying either an NFκB response element or a constitutive SFFV promoter driving luciferase. Here for the first time we report the generation of light emitting somatic transgenic rodents with a wider spread of transgene expression, following a single neonatal intravenous or intracranial administration of AAV8 biosensors. We validated the biosensing technology by administering LPS to model systemic inflammation and showed a significant increase in light emission. This technology will allow for expedited investigations regarding signalling pathways activated in disease processes and complement existing germline transgenic light producing technology by maximising the use, and reducing the numbers of animals used in biomedical research.

## Results

### Neonatal intravenous administration of AAV8-CMV-GFP vector

We sought to determine the bio-distribution of GFP expression following neonatal (Post-neonatal day 1, P1) intravenous administration of AAV8-CMV-eGFP. On the day of birth, mice received 40μl of vector (1 × 10^13^ vector genomes/ml) via the superficial temporal vein. One month later, mice were harvested and GFP expression was analysed, using a DFC420 microscope.

Prior to gross dissection and further fixation, skin was removed and mice were viewed under a stereoscopic fluorescence microscope. Strong, extensive and widespread GFP expression was observed throughout the body, although the highest levels of expression were most noticeable in the musculature (Figure 1). Strong transduction was observed within the heart (B), liver (C), kidney (D), skeletal muscle (E), eye (F), brain (G), and myenteric plexus (H).

**Figure 1.**
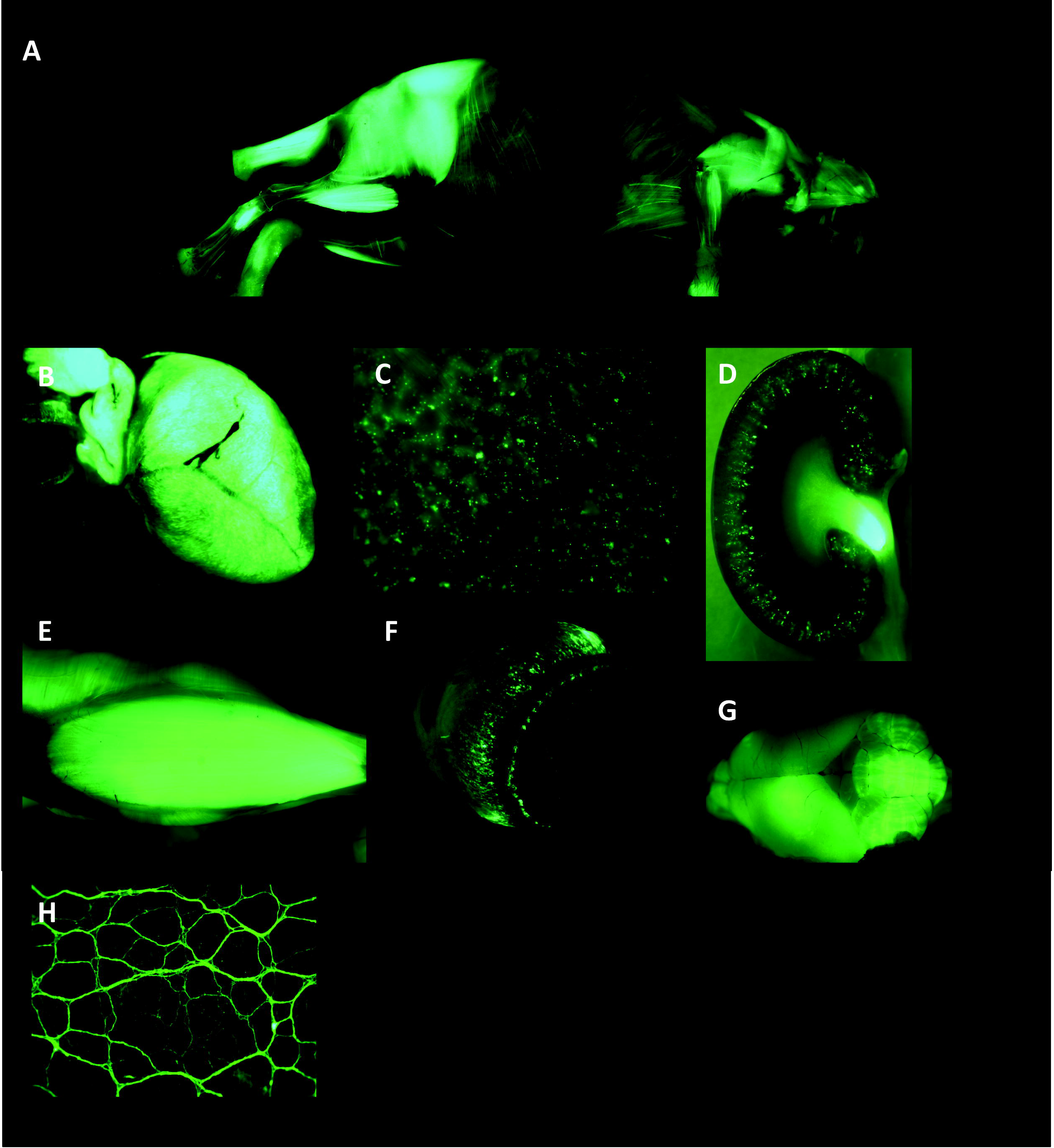
Systemic distribution of GFP after a single neonatal intravenous administration of AAV8-CMV-GFP. New-born mice received an intravenous administration of AAV8-CMV-GFP vector (n=3). At 1 month of age the mice were harvested and the *ex vivo* analysis of GFP expression revealed widespread systemic distribution (A). Strong GFP expression was observed within the heart (B), liver (C), kidney (D), muscle (E), eye ball (F), brain (G) and the myenteric plexus (H).

In order to assess the expression profile in the CNS, the brains from injected and non-injected control mice were sectioned and immunohistochemistry was conducted for GFP expression. This revealed extensive and widespread GFP expression (Supplementary figure 1). Further examination under higher magnification of discrete areas of the brain including the primary motor cortex, the somatosensory barrel field (S1BF), piriform cortex, dentate gyrus, cerebellum, and the gigantocellular nucleus revealed transduction of cells with both neuronal and glial morphology (Supplementary figure 1).

Further to this we investigated whether AAV vector or GFP transgene expression triggers an inflammatory response after neonatal intracranial injections. Microglia activation was examined in all brains and compared to brain tissue from Ppt1^−/−^ (palmitoyl protein thioesterase 1) mice which are a known model of microglia activation^18^. Extensive microglia engorgement and activation was observed in the Ppt1^−/−^ mice and no noticeable activation of microglia was observed in the non-injected and AAV8 injected brains (Supplementary figure 2).

**Supplementary figure 1 - Immunohistochemical detection and quantification of GFP expression 1 month post intravenous injection of AAV8 into PI neonates.** PI neonates were intravenously administered with self-complementary AAV8-CMV-eGFP (n=3). Uninjected mice were used as negative controls (n=3). At P30 the mice were culled and the brains were removed for sectioning and immunohistochemical staining using anti-GFP antibody and DAB staining. The sections were examined by light microscopy and representative images were taken from the pre-frontal cortex, striatum, hippocampus, piriform lobes and the cerebellum (A). Various discrete areas of the brains were examined under higher magnification including the primary motor cortex, S1BF, piriform cortex, dentate gyrus, cerebellum and gigantocellular nucleus (B). Quantitative measurement of staining in discrete areas of the brain was conducted by thresholding analysis in the S1BF, caudate putamen (CPu), CA1 region of the hippocampus (CA1), piriform cortex and the cerebellar nodule (lOcb). The data was plotted as the mean ± S.D.

**Supplementary figure 2. Immunohistochemical and quantitative analysis of microglia activation.** Brain sections from both injected and uninjected mice were analysed for microglia-mediated immune response. The sections were probed using antibodies against the microglia marker CD68 and detected using DAB. As a positive control, brain sections taken from Ppt1^−/−^ knockout mice (n=3) were included since they have a known microglia immune response. Representative high magnification images were taken of stained microglia from Ppt1^−/−^ mice, uninjected mice and AAV8 administered mice (A). Quantitative measurement of staining intensity was conducted by thresholding analysis (B). The data is plotted as the mean ± SEM (n=3).

### Production of AAV8 biosensors

An AAV8 producer plasmid was created containing a Gateway^®^ accepter site (Invitrogen). The Gateway^®^ sequence was cloned into the backbone and was placed upstream of a minimal promoter driving a codon-optimised luciferase transgene and an enhanced GFP linked by a bicistronic linker, T2A (Supplementary figure 3). We have now assembled an extensive library of transcription factor binding elements in pENTR shuttle plasmids and these are shown in supplementary figure 4. We selected the NFκB response element and an SFFV viral promoter for the insertion into the AAV gateway backbone. These two were chosen as they have been validated by both *in vitro* and *in vivo* means in our lentiviral system^3^.

AAV8 biosensor vectors were generated using the AAV8-SFFV-Luc-T2A-eGFP and AAV8-NFκB-Luc-T2A-eGFP backbones.

**Supplementary figure 3 – Plasmid configuration for AAV8-GW-Luc-T2A-GFP.** Between the AAV2 ITR sequences is the Gateway cloning site, followed by a 3xFlag tag upstream of the codon-optimised luciferase transgene. A bicistronic linker, T2A sequence, is placed upstream of an enhanced GFP. Gene expression is enhanced with the presence of woodchuck hepatitis virus post-transcriptional regulatory element (WPRE).

**Supplementary figure 4 – Table of response element cloned into the lentiviral biosensor.** A number of response elements are cloned into the lentiviral backbone using Gateway^®^ cloning. Each response element has been validated *in vitro* with the appropriate agonist.

### Neonatal administration of AAV8 biosensors

Having observed widespread transgene expression after a single neonatal administration of an AAV8-CMV-GFP vector, we chose to investigate the NFκB signalling expression profile by neonatal injection of the AAV8-NFκB-LUC-2A-GFP biosensor. We selected AAV8-SFFV-Luc-2A-GFP as a constitutively expressed control and to allow comparison with previous experiments using lentivirus vectors^3^.

At P1 of development, mice received a 30μl intravenous (IV) administration of AAV8 SFFV or AAV8 NFκB biosensor (l×l0^13^ vg/ml). Mice underwent whole-body bioluminescence imaging over the course of development to quantitate luciferase expression.

Following IV injection of the AAV8 NFκB biosensor, luciferase expression was strongest in the spine, thorax, paws, lower abdominal and the mouth (Figure 2A). In contrast, IV injection of the AAV8 SFFV biosensor resulted in whole-body luciferase expression but with strongest expression in the lower abdomen (Figure 2A). The luciferase expression was quantified and showed stable transgene expression over development (Figure 2B and C).

**Figure 2.**
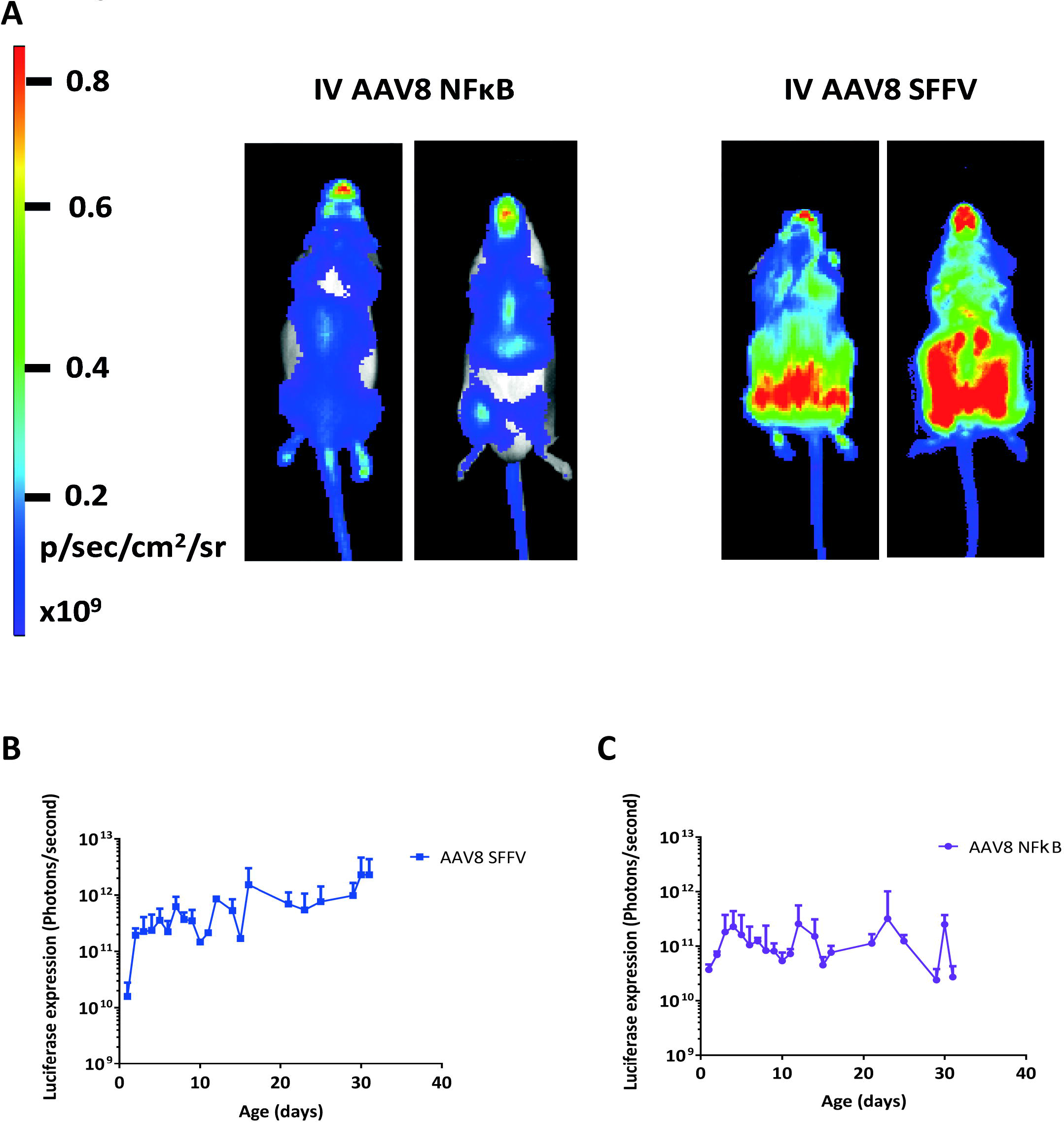
Whole-body bioluminescence imaging of mice which have received an intravenous administration of AAV8 SFFV or AAV8 NFκB vectors. Newborn mice received intravenous injections of either AAV8 SFFV or AAV8 NFκB biosensors (n = 6 per group). We observed expression profiles from the two different AAV8 biosensors (A), the same mouse was imaged on its front and back. Luciferase expression was quantified by whole-body bioluminescence imaging for a month (B and C), (mean +/− SD).

Additionally, to restrict the expression profile within the CNS and PNS, intracranial (IC) injections were also performed with AAV8 SFFV and AAV8 NFκB biosensors. P1 pups received 5μl (1×10^13^ vg/ml) of either biosensor, and whole-body bioluminescence imaging was undertaken over the course of development. Luciferase expression from the IC injected AAV8 N FKB biosensors was similar to that seen in the IV injected mice, predominantly in the spine, thorax, paws, mouth, eyes and tail (Supplementary figure 5A).

A month after IC or IV injection of AAV8 SFFV or NFκB biosensor, the mice were harvested and immunohistochemistry performed to detect GFP expression in the brain. After IC injection of AAV8 SFFV or NFκB biosensor, GFP expression was predominantly neuronal within the dentate gyrus and CA1 and CA3 regions of the hippocampus (Supplementary Figure 6A and C). However, following IV AAV8 SFFV, a mixture of neurons and astrocytes were found to be positive for GFP within the hippocampal region (Supplementary Figure 6B). After IV injection of AAV8 NFκB, a low level of GFP expression was found within a small number of astrocytes and neuronal cells (Supplementary Figure 6D).

Further to this, on a separate cohort of intravenously injected mice, we were able to show extensive whole-body bioluminescence at 240 days of development (Figure 3).

**Figure 3.**
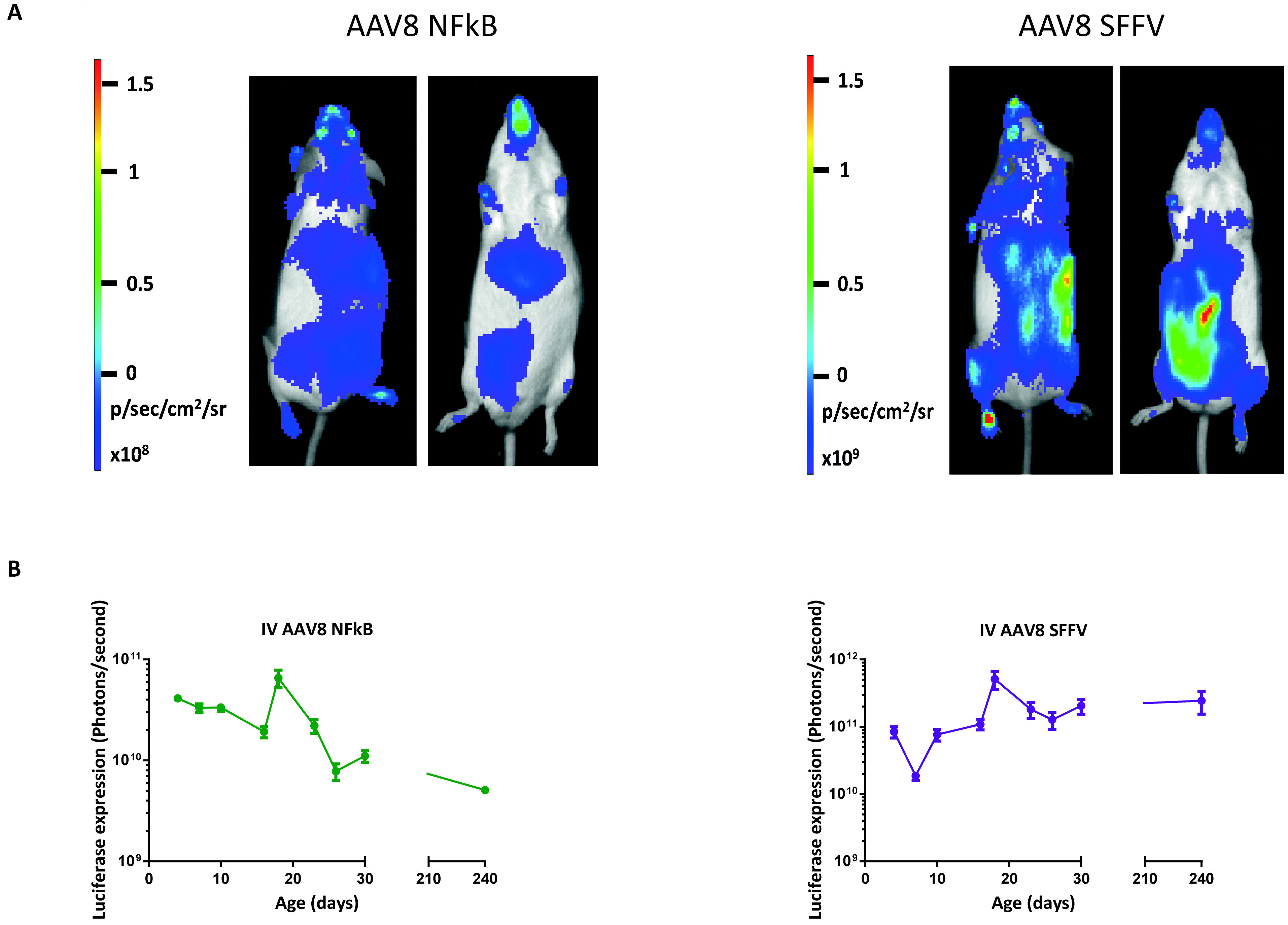
Whole-body bioluminescence imaging of mice which have received an intravenous administration of AAV8 NFκB vectors up to 240 days of development. Newborn mice received intravenous injections of AAV8 NFκB biosensors (n = 8 per group). (A) the same mouse was imaged on its front and back. Luciferase expression was quantified by whole-body bioluminescence imaging for 8 months (B), (mean +/− SD).

**Supplementary figure 5 – Luciferase expression and bio-distribution following intracranial injections of AAV8 NFκB or AAV8 SFFV vectors.** Mice received an intracranial administration of AAV8 NFκB or SFFV vectors (n = 6 per group) on the day of birth, PI. The mice underwent whole-body bioluminescence imaging over development up to day 31 of development. The bio-distribution of luciferase expression from the two biosensors differed (A), the same mouse was imaged on its front and back. Luciferase expression was quantified for a month (B and C) (mean +/− SD).

**Supplementary figure 6 – Neuronal and astrocytic targeting after intravenous or intracranial administration of AAV8 SFFV or NFκB biosensors.** Brain sections from injected mice were used to determine GFP positive cells. IC AAV8 SFFV revealed a predominant neuronal transduction within the dentate gyrus and the CA1 and CA3 regions of the hippocampus (A). IV AAV8 SFFV sections showed a mixture of astrocytic and neuronal cell morphology (B). 1C AAV8 NFκB targeted neuronal cells (C) whereas IV AAV8 NFκB transduced astrocytes and neurons retrospectively (D).

### Validation of the AAV8 NFκB biosensor

To assess whether the AAV8 NFκB could be exploited to report on NFκB signalling in pathological states, mice received ultra-pure lipopolysaccharide (LPS) which only acts through toll-like receptor 4 to induce translocation of NFκB from the cytosol to the nucleus^19^. At day 132 after AAV administration, a single dose of LPS was administered intraperitoneally to both IV and 1C injected groups. Bioimaging was taken at 4, 24, 48, 72 and 96 hours before (to correct for any perturbance in the signal caused by multiple imaging) and after LPS administration. A single administration of LPS resulted in a significant increase in luciferase expression in IV (Figure 4; p < 0.0001) and 1C AAV8 NFκB mice (Supplementary figure 7; p = 0.006).

**Figure 4.**
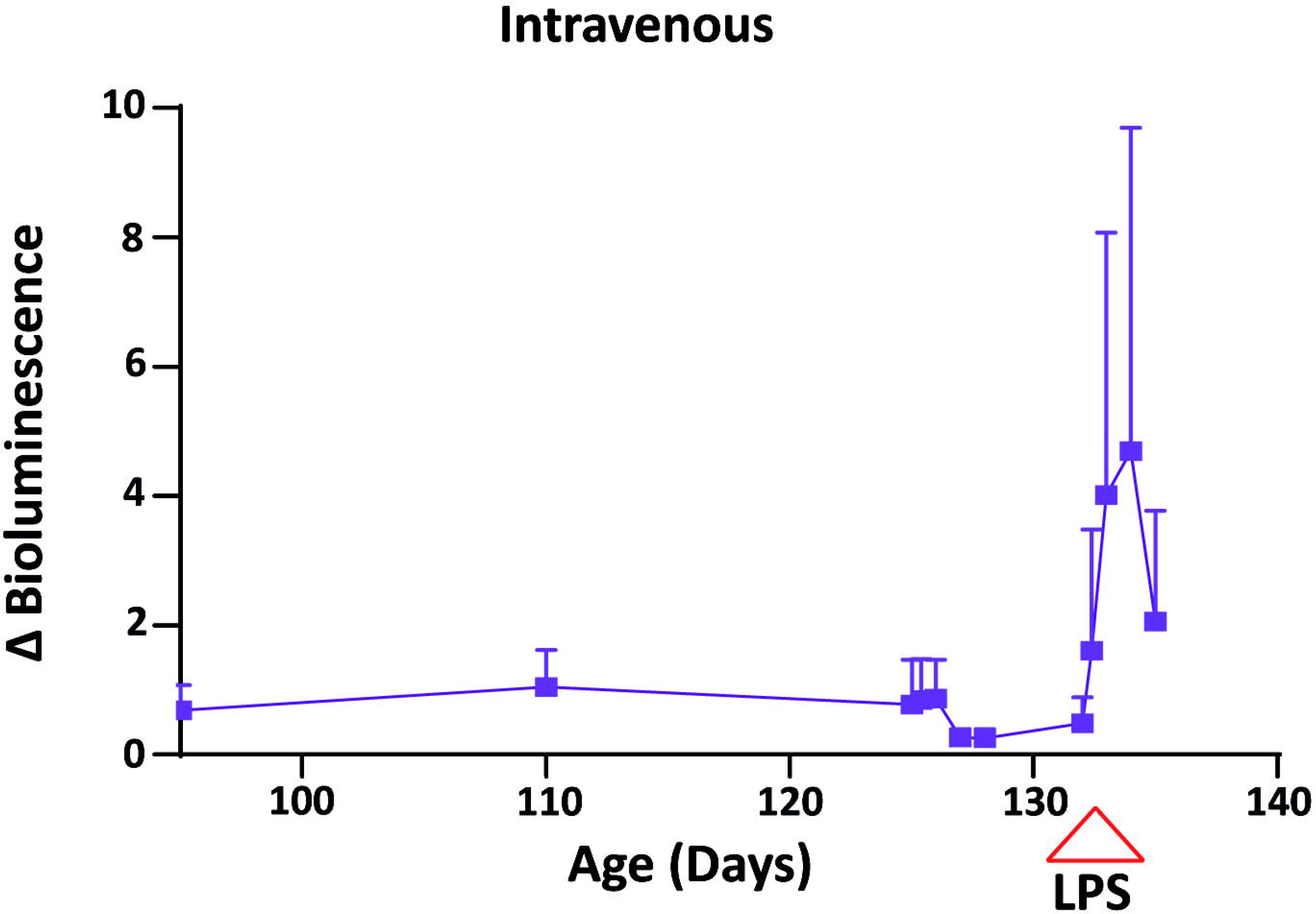
Up-regulation of luciferase signal mediated by a single dose of LPS. Bioluminescence imaging persisted more than 4 months over development in mice which received intravenous administration of the AAV8 NFκB biosensor. At day 132 of development all the mice received a single dose of LPS which resulted in a significant up-regulation in luciferase expression in the IV injected mice, p < 0.0001.

**Supplementary figure 7 – Up-regulation of luciferase signal mediated by a single dose of LPS in intracranially injected mice.** Bioluminescence imaging persisted more than 4 months over development in mice which received an intracranial administration of AAV8 NFκB biosensor. At day 132 of development all the mice received a single dose of LPS which resulted in a significant up-regulation in luciferase expression in 1C injected mice, p = 0.006.

## Discussion

In this study, we have shown for the first time that light producing somatic-transgenic rodents can be produced by a single administration of an AAV8 biosensor. Specifically, we have demonstrated that AAV8 vectors cross the blood brain barrier (BBB) and mediate systemic transgene expression after a single neonatal intravenous or intracranial administration.

Our results showed a systemic spread of transgene expression by a neonatal intravenous administration of a self-complementary AAV8 vector. This data agrees with previous work conducted by Nakai et al. which demonstrated that AAV8 vectors cross the BBB and result in transduction of both neuronal and glial cells in the brain, albeit with a low efficiency^20^. They also presented transduction of cells within the liver, skeletal muscle, smooth muscle, cardiac muscle and pancreas^20^. However, we were able to show a much more extensive spread of transgene expression within the CNS and visceral organs such as the kidney and large intestines. The strong and widespread GFP expression observed in our study maybe due to a number of factors including the use of a different promoter and route of administration.

Our analysis also demonstrated an improved safety profile within the CNS after a neonatal intravenous administration. It has been reported that lentiviral vector mediated expression of GFP elicits toxicity in Purkinje cells of the developing mouse cerebellum. This toxicity was attributed mainly to GFP overexpression although the authors could not discount the lentiviral vectors *per se*, or genotoxicity caused by vector integration in neuronal DNA as aggregating factors^21^. In contrast, we detected no microglial activation in the CNS suggesting that inflammation or gliosis caused by the injection, the presence of vector, or expression of both luciferase and GFP did not occur. This suggests that the AAV system we describe may be preferable to our previously described lentiviral vector system^2^.

We preformed neonatal intracranial injections of our AAV8 biosensors to monitor whether we could restrict the expression profile to the CNS and PNS. However, our results revealed that the intracranial injected mice showed a similar expression profile to the mice which received an intravenous administration. As previous work has shown that after a single adult intravenous injections of AAV8, transgene expression was observed within the CNS and also within the cervical spinal cord, thoracic spinal cord, and lumbar spinal cord sections. However, the expression profile was not significant compared to other serotypes such as AAVrh.8^22^

In order to overcome this wide distribution profile, future experiments would involve the assessment of additional AAV capsids. These would include AAV-PHP.B, which is a new variant AAV capsid that transduces the CNS 40-fold greater than AAV9 after a single adult intravenous administration and targets astrocytes and neurons^23^. Furthermore, AAV-PHP.eB an enhanced variant of AAV-PHP.B, efficiently transduces the CNS and PNS greater than the AAV-PHP.B and predominately targets neurons^24^. Therefore, in order to achieve CNS restricted and cell specific transduction profiles different configurations of AAV capsids would have to be used.

We have established a unique technology which allows the production of somatic-transgenic light emitting rodents. We have synthesised a library of >20 response elements, >10 promoters, most of which we have validated in the context of a lentiviral backbone *in vitro*, and several of them *in vivo*. To exploit the advantages of AAV, we gene synthesised an AAV backbone containing a Gateway acceptor site, and strategically placed multi-cloning sites with unique restriction sites to permit easy cloning.

As proof-of principle we inserted the NFκB response element, consisting of octuplet cognate binding elements, into the AAV backbone. A potential concern with introducing such sequences into the viral backbone is the loss of tandem repeats following propagation and amplification in bacteria^25^. However, sequencing of large-scale DNA preparations of this backbone confirmed that the repeats remained intact.

Here we demonstrate that administration of AAV biosensors by a single neonatal intravenous administration resulted in widespread luciferase expression, in comparison to our lentiviral biosensors which only transduced the liver^3^. The bio-distribution of luciferase expression differed between the constitutive AAV8 SFFV biosensor and the NFκB biosensor. A previous study demonstrated NFκB biosensor activity following administration of AAV by pancreatic infusion in adult mice^26^. Baseline expression rose rapidly in those mice but fell to modest levels by 1 month. This may be attributable to an anti-luciferase or anti-capsid cytotoxic T-cell response eliminating transduced cells. In contrast, we observed stable expression for up to at least five months; it has been shown that neonatal gene transfer administration results in immune tolerization to the transgenic protein when using retrovirus^27^, AAV^28^ and adenovirus^29^ vectors.

Here we present for the first time the generation of somatic-transgenic mice with the use of AAV viral vectors. We have also shown widespread transgene expression after a single intracranial or intravenous neonatal administration of the AAV biosensors. We have validated the AAV biosensors with the use of agonists. This technology not only complements existing germline transgenic rodents but also maximises the use and reduces the numbers of animals used in biomedical research^30^. We have generated a non-invasive gene marking technology, which can be applied to systemic disease models. With the use of Gateway^®^ cloning the assessment of different response elements can easily be achieved.

## Methods and materials

### Generation of the AAV biosensor plasmids

The AAV8-Gateway^®^-Luc-T2A-eGFP was synthesised by Aldevron (Aldevron, North Dakota, USA). This AAV plasmid consisted of a Gateway^®^ sequence, placed upstream of a codon-optimised luciferase transgene, linked to an enhanced GFP by a bicistronic linker, T2A. The response element NFκB and constitutive promoter SFFV was cloned into an pENT plasmid^3^. By using the Gateway^®^ cloning kit (Invitrogen, Manchester, UK) and using the manufactures guidelines, the response element NFκB and constitutive promoter SFFV were individually inserted into the AAV backbone to generate the following plasmids; pAAV8-SFFV-Luc-T2A-eGFP and pAAV8-NFκB-Luc-T2A-eGFP.

### Recombinant AAV production

Recombinant AAV was produced, purified and titered using standard procedures. Briefly, HEK293T cells were double transfected with the pAAV8-SFFV-Luc-2A-eGFP or pAAV8-NFκB-Luc-2A-eGFP plasmid, and the pDG8 plasmid expressing AAV2 Rep, AAV8 Cap gene and adenovirus 5 helper functions (Plasmid Factory, Bielefeld, Germany) using polyethylenimine (PEImax, Polysciences Inc). After 72 hours of incubation at 37°C, the cells were harvested by centrifugation and then lysed by freeze-thawing in lysis buffer. In parallel, the virus-containing supernatant was harvested and precipitated by using ammonium sulphate salt. Cell lysate and precipitated supernatant were treated with benzonase, clarified by centrifugation and filtered at 0.22μm before purification.

The recombinant AAV virus preparations were purified by iodixanol step gradient: the viral preparation was overlaid with increasing concentrations of iodixanol (15%, 25%, 40% and 60%, OptiPrep; Sigma-Aldrich, Dorset, UK). The tubes were centrifuged at 40,000 rpm for 3 hours at 18°C in a Sorvall Discovery 90SE ultracentrifuge using a TH641 rotor (Thermo Scientific, Paisley, UK). The vector was extracted from the 40% fraction with a 19-gauge needle. Purified vector fractions were dialysed against PBS overnight.

Real-time PCR and alkaline gel electrophoresis were used to assess the viral genome titers and integrity of the viral genome^12,13^, the capsid titers were determined by SDS PAGE electrophoresis^14^.

AAV8 containing the enhanced GFP gene driven by the cytomegalovirus promoter (AAV8-CMV-eGFP) in a self-complementary configuration was obtained from the UPenn Vector Core facility (Philadelphia, USA) at a titre of 1×10^13^ vg/ml.

### Animal procedures

Outbred CD1 mice and MF1 mice used in this study were supplied by Charles Rivers Laboratories. All procedures were performed under United Kingdom Home Office Project License 70/8030, approved by the ethical review committee and followed institutional guidelines at University College London.

### Neonatal Intracranial and Intravenous injections

For intracranial injections, mice (random mix of males and females) were subjected to brief hypothermic anaesthesia on the day of birth, followed by unilateral injections of concentrated AAV vector (5μl in total; 1 × 10^13^ vector genomes/ml) into the cerebral lateral ventricles using a 33 gauge Hamilton needle (Fisher Scientific, Loughborough, UK), following co-ordinates provided by Kim et al.^15^. For intravenous injections, pups were subjected to brief hypothermic anaesthesia followed by intravenous injections of AAV vectors into the superficial temporal vein^16^, with a total volume of 25μl (1 × 10^13^ vg/ml). The neonates were then allowed to return to normal temperature before placing them back with the dam.

### Whole-body bioluminescence imaging

Where appropriate, mice were anaesthetised with isoflurane with 21% oxygen (Abbotts Laboratories, London, UK). D-luciferin (Gold Biotechnologies, ST Louis, USA) was administered by intraperitoneal injection at a concentration of 15mg/mL. Mice were imaged 5 minutes after luciferin injection using a cooled charged-coupled device camera, (IVIS Lumina II, Perkin Elmer, Coventry, UK) for between 1 second and 5 minutes. Photon output of the whole-body was measured using Living Image Software (Perkin Elmer) and light output quantified and expressed as photons per second per centimetre squared per steradian (photons/second/cm^2^/sr).

### Collection of brain tissues

Mice were anaesthetised at day 35 of development using Isoflurane and the right cardiac atrium was incised, followed by injection of heparinized PBS into the left cardiac ventricle. The brains were removed and fixed in 4% paraformaldehyde (PFA) and then cryoprotected in 30% sucrose in 50mM PBS. Brains were sectioned using a sliding microtome (Carl Zeiss, Welwyn Garden City, UK) to generate 40[im transverse sections^16^. Sections were stored in 30% sucrose in TBS, ethylene glycol and 10% sodium azide.

### Immunoperoxidase immunohistochemistry

To visualise CD68 and GFP immunoreactivity, sections were treated with 30% H_2_O_2_ in TBS for 30 minutes. They were blocked with 15% rabbit serum for CD68 (Vector Laboratories, Cambridge, UK) and goat serum for GFP (Vector Laboratories) in Tris buffered saline and Tween 20 (TBST) for 30 minutes. This was followed by the addition of primary antibody, rat anti-mouse CD68 (1:100; Biorad, Hertfordshire, UK), mouse anti-GFP (1:10,000; Abcam, Cambridge, UK) in 10% serum and TBST and left on a gentle shaker over night at 4°C. The following day the sections were treated with the secondary rat anti-rabbit antibody for CD68 (1:1000; Vector Laboratories) and goat anti-rabbit (1:1000 dilution; Vector Laboratories) in 10% rat or goat serum in TBST for 2 hours. The sections were incubated for a further 2 hours with Vectastain ABC (Vector Laboratories). 0.05% of 3,3’-diaminobenzidine (DAB) was added and left for a couple of minutes. Sections were transferred to ice cold TBS. Individual brain sections were mounted onto chrome gelatine-coated Superfrost-plus slides (VWR, Poole, UK) and left to dry for 24 hours. The slides were dehydrated in 100% ethanol and placed in Histoclear (National Diagnostics, Yorkshire, UK) for 5 minutes before adding a coverslip with DPX mounting medium (VWR). DAB stained sections in supplementary figures 1 and 2 were viewed using an Axioskop 2 Mot microscope (Carl Zeiss Ltd.) and representative images were captured using an Axiocam HR camera and Axiovision 4.2 software (Carl Zeiss Ltd.). DAB stained sections in supplementary figure 7 were viewed using Leica MZ16F microscope software.

### Quantitative analysis of immunohistological staining

GFP and CD68 expression was quantified by thresholding analysis as previously described^17,18^. Briefly, 40 non-overlapping RGB images were taken from four consecutive sections through the somatosensory barrel field (S1BF), caudate putamen (Cpu), the *Cornu Ammonis* region 1 of the hippocampus, (CA1), piriform cortex (piriform cort), and the lOCb region of the cerebellum (10Cb). The images were captured using a live video camera (JVC, 3CCD, KY-F55B) mounted onto a Zeiss Axioplan microscope using the ×40 objective lens. All camera and microscope calibration and settings were kept constant throughout the image capture period. Images were analysed for optimal segmentation and immunoreactive profiles were determined using Image-Pro Plus (Media Cybernetics). Foreground immunostaining was accurately defined according to averaging of the highest and lowest immunoreactivities within the sample population for a given immunohistochemical marker (per colour/filter channel selected) and measured on a scale from 0 (100% transmitted light) to 255 (0% transmitted light) for each pixel. This threshold setting was constant to all subsequent images analyzed for the antigen used. Immunoreactive profiles were discriminated in this manner to determine the specific immunoreactive area (the mean grey value obtained by subtracting the total mean grey value from non-immunoreacted value per defined field). Macros were recorded to transfer the data to a spreadsheet for subsequent statistical analysis.

### Statistics

The data from supplementary figures 1 and 2 were plotted graphically as the mean percentage area of immune-reactivity per field ± S.D or S.E.M. for each region. For measuring the fold change of bioluminescence (Figure 4 and supplementary figure 8), a median of luciferase expression from days 48 to 84 was taken and the raw luciferase values were divided by the median. The median was used, as it is a much more robust measure of central tendency. Two-way ANOVA with and a Sidak’s multiple comparison was performed in Figure 4 and supplementary figure 8.

## Acknowledgements

RK and SNW received funding from MRC grants MR/P026494/1 and MR/R015325/1, and from SPARKS grant 17UCL01. AAR and SNW received funding from UK MRC grant MR/N026101/1. TRM, AAR and SNW received part funding from UK NC3Rs grant NC/L001780/1. SNW, DPP, TRM and SMKB received funding from ERC grant “Somabio” 260862. DPP received funding from UK MRC grant MR/N019075/1. NS received a Clinical Research Training Fellowship from Wellbeing of Women. JAD is supported by CONICYT Becas Chile Doctoral Fellowship program 72160294. JRC received funding from NIHR GOSH BRC grant 17BX23. This research was supported by the NIHR Great Ormond Street Hospital Biomedical Research Centre. The views expressed are those of the authors and not necessarily those of the NHS, the NIHR or the Department of Health. EH received funding from MRC grant MR/N022890/1. JDC and AMSW was supported by The Wellcome Trust (GR079491MA), Batten Disease Family Association, and the Batten Disease Support and Research Association.

## Author contributions

SNW, AAR, RK, SMKB and TRM developed experimental design. RK, AAR, AMSW, NS, JAD, DPP, NPM, MH, JMKMD, JRC, JDC, EH, TRM, SMKB and SNW contributed to data. RK, AAR, AMSW, JRC, EH, TRM, SMKB and SNW contributed to manuscript.

## Additional information

**Supplementary information** – accompanies this paper.

**Competing financial interests** – authors declare no competing financial interests.

## References

1. Delhove, J. M. K. M. et al. Longitudinal in vivo bioimaging of hepatocyte transcription factor activity following cholestatic liver injury in mice. Sci. Rep. 7, 41874 (2017).

2. Karda, R. et al. Continual conscious bioluminescent imaging in freely moving somatotransgenic mice. Sci. Rep. 7, 6374 (2017).

3. Buckley, S. M. K. et al. In vivo bioimaging with tissue-specific transcription factor activated luciferase reporters. Sci. Rep. 5, 11842 (2015).

4. Hawkins, K. E. et al. NRF2 Orchestrates the Metabolic Shift during Induced Pluripotent Stem Cell Reprogramming. Cell Rep. 14, 1883–1891 (2016).

5. Teasdale, J. E. et al. Cigarette smoke extract profoundly suppresses TNFα-mediated proinflammatory gene expression through upregulation of ATF3 in human coronary artery endothelial cells. Sci. Rep. 7, 39945 (2017).

6. Ivacik, D., Ely, A., Ferry, N. & Arbuthnot, P. Sustained inhibition of hepatitis B virus replication in vivo using RNAi-activating lentiviruses. Gene Ther 22, 163–171 (2015).

7. Carlon, M. S. et al. Immunological Ignorance Allows Long-Term Gene Expression After Perinatal Recombinant Adeno-Associated Virus-Mediated Gene Transfer to Murine Airways. Hum. Gene Ther. 12, 1–12 (2014).

8. Zincarelli, C., Soltys, S., Rengo, G. & Rabinowitz, J. E. Analysis of AAV serotypes 1-9 mediated gene expression and tropism in mice after systemic injection. Mol. Ther. 16, 1073–1080 (2008).

9. Inagaki, K. et al. Robust systemic transduction with AAV9 vectors in mice: efficient global cardiac gene transfer superior to that of AAV8. Mol. Ther. 14, 45–53 (2006).

10. Wang, Z. et al. Adeno-associated virus serotype 8 efficiently delivers genes to muscle and heart. Nat. Biotechnol. 23, 321–328 (2005).

11. Foust, K. D., Poirier, A., Pacak, C. a, Mandel, R. J. & Flotte, T. R. Neonatal intraperitoneal or intravenous injections of recombinant adeno-associated virus type 8 transduce dorsal root ganglia and lower motor neurons. Hum. Gene Ther. 19, 61–70 (2008).

12. Zeltner, N., Kohlbrenner, E., Clément, N., Weber, T. & Linden, R. M. NIH Public Access. Gene 17, 872–879 (2011).

13. Fagone, P. et al. Systemic Errors in Quantitative Polymerase Chain Reaction Titration of Self-Complementary Adeno-Associated Viral Vectors and Improved Alternative Methods. Hum. Gene Ther. Methods 23, 1–7 (2012).

14. Kohlbrenner, E. et al. Quantification of AAV Particle Titers by Infrared Fluorescence Scanning of Coomassie-Stained Sodium Dodecyl Sulfate-Polyacrylamide Gels. Hum. Gene Ther. Methods 23, 198–203 (2012).

15. Kim, J. Y. et al. Viral transduction of the neonatal brain delivers controllable genetic mosaicism for visualising and manipulating neuronal circuits in vivo. Eur. J. Neurosci. 37, 1203–1220 (2013).

16. Rahim, A. et al. Intravenous administration of AAV2/9 to the fetal and neonatal mouse leads to differential targeting of CNS cell types and extensive transduction of the nervous system. 3505–3518 (2011). doi:10.1096/fj.11-182311

17. Bible, E., Gupta, P., Hofmann, S. L. & Cooper, J. D. Regional and cellular neuropathology in the palmitoyl protein thioesterase-1 null mutant mouse model of infantile neuronal ceroid lipofuscinosis. Neurobiol. Dis. 16, 346–359 (2004).

18. Kielar, C. et al. Successive neuron loss in the thalamus and cortex in a mouse model of infantile neuronal ceroid lipofuscinosis. Neurobiol. Dis. 25, 150–162 (2007).

19. Hirschfeld M, Ma Y, Weis J, Vogel S, W. J. Cutting edge: repurification of lipopolysaccharide eliminates signaling through both human and murine toll-like receptor 2. J Immunol Ref. 165, 618–622 (2000).

20. Nakai, H. et al. Unrestricted hepatocyte transduction with adeno-associated virus serotype 8 vectors in mice. J. Virol. 79, 214–24 (2005).

21. Sawada, Y. et al. High transgene expression by lentiviral vectors causes maldevelopment of Purkinje cells in vivo. Cerebellum 9, 291–302 (2010).

22. Yang, B. et al. Global CNS Transduction of Adult Mice by Intravenously Delivered rAAVrh.8 and rAAVrh.10 and Nonhuman Primates by rAAVrh.10. Mol. Ther. 22, 1299–309 (2014).

23. Deverman, B. E. et al. Cre-dependent selection yields AAV variants for widespread gene transfer to the adult brain. Nat. Biotechnol. advance on, 1–7 (2016).

24. Chan, K. Y. et al. Engineered AAVs for efficient noninvasive gene delivery to the central and peripheral nervous systems. Nat. Neurosci. 20, 1172–1179 (2017).

25. Bzymek, M. & Lovett, S. T. Instability of repetitive DNA sequences: The role of replication in multiple mechanisms. Proc. Natl. Acad. Sci. 98, 8319–8325 (2001).

26. Orabi, A. I. et al. Dynamic imaging of pancreatic NF-κB activation in live mice using AAV infusion and bioluminescence. J. Biol. Chem. 291, jbc. M115. 647933 (2015).

27. Zhang, J., Xu, L., Haskins, M. E. & Ponder, K. P. Neonatal gene transfer with a retroviral vector results in tolerance to human factor IX in mice and dogs. Blood 103, 143–151 (2004).

28. Shi, Y., Falahati, R., Zhang, J., Flebbe-Rehwaldt, L. & Gaensler, K. M. L. Role of antigen-specific regulatory CD4+CD25+ T cells in tolerance induction after neonatal IP administration of AAV-hF.IX. Gene Ther. 20, 987–996 (2013).

29. Nivsarkar, M. S. et al. Evidence for contribution of CD4+ CD25+ regulatory T cells in maintaining immune tolerance to human factor IX following perinatal adenovirus vector delivery. J. Immunol. Res. 2015, 397879 (2015).

30. Knight, K. Implementing the 3Rs: improving experimental approaches in animal biology. J. Exp. Biol. 219, 2414–2415 (2016).

